# Novel TRPM7 inhibitors with potent anti-inflammatory effects *in vivo*

**DOI:** 10.1101/2023.05.22.541802

**Authors:** Gregory W. Busey, Mohan C. Manjegowda, Tao Huang, Wesley H. Iobst, Shardul S. Naphade, Joel A. Kennedy, Catherine A. Doyle, Philip V. Seegren, Kevin R. Lynch, Bimal N. Desai

**Affiliations:** Pharmacology Department, University of Virginia School of Medicine, Pinn Hall, 1340 Jefferson Park Avenue, Charlottesville, VA 22908. USA; Carter Immunology Center, University of Virginia School of Medicine, 345 Crispell Dr. MR-6, Charlottesville, VA 22908. USA; University of Virginia Cardiovascular Research Center, 415 Lane Rd, Charlottesville, VA 22908. USA

**Keywords:** TRPM7, inflammation, sepsis, anti-inflammatory, drug development, ion channel, neuroinflammation, immunopharmacology, lipopolysaccharide, small-molecule inhibitor

## Abstract

TRPM7, a TRP channel with ion conductance and kinase activities, has emerged as an attractive drug target for immunomodulation. Reverse genetics and cell biological studies have already established a key role for TRPM7 in the inflammatory activation of macrophages. Advancing TRPM7 as a viable molecular target for immunomodulation requires selective TRPM7 inhibitors with *in vivo* tolerability and efficacy. Such inhibitors have the potential to interdict inflammatory cascades mediated by systemic and tissue-specialized macrophages. FTY720, an FDA-approved drug for multiple sclerosis inhibits TRPM7. However, FTY720 is a prodrug and its metabolite, FTY720-phosphate, is a potent agonist of sphingosine 1-phosphate (S1P) receptors. In this study, we tested non-phosphorylatable FTY720 analogs, which are inert against S1PRs and well tolerated *in vivo*, for activity against TRPM7 and tissue bioavailability. Using patch clamp electrophysiology, we show that VPC01091.4 and AAL-149 block TRPM7 current at low micromolar concentrations. In culture, they act directly on macrophages to blunt LPS-induced inflammatory cytokine expression, an effect that is predominantly but not solely mediated by TRPM7. We found that VPC01091.4 has significant and rapid accumulation in the brain and lungs, along with direct anti-inflammatory action on alveolar macrophages and microglia. Finally, using a mouse model of endotoxemia, we show VPC01091.4 to be an efficacious anti-inflammatory agent that arrests systemic inflammation *in vivo*. Together, these findings identify novel small molecule inhibitors that allow TRPM7 channel inhibition independent of S1P receptor targeting. These inhibitors exhibit potent anti-inflammatory properties that are mediated by TRPM7 and likely other molecular targets that remain to be identified.

## Introduction

Transient receptor potential melastatin-like 7 (TRPM7) is a member of the TRP superfamily of ion channels, a major class of drug targets (Koivisto et al., 2022, Moran, 2018). TRPM7 is one of two human *chanzymes* encoding both a non-selective cation channel and an intracellular serine/threonine kinase domain (Nadler et al., 2001, Runnels et al., 2001). *In utero*, TRPM7 is broadly expressed in all tissues and is essential for embryogenesis. After embryogenesis, its expression subsides to low levels in most tissues but remains at relatively high levels in hematopoietic cells (Jin et al., 2008). In adult mice, post-embryonic deletion of *Trpm7* in various organs is well-tolerated after the completion of organogenesis (Jin et al., 2012) making TRPM7 a potential drug target for niche immunopharmacology. Since TRPM7 currents (I_TRPM7_ or simply, I_M7_) are readily detectable in immune cells, we explored and established significant functions of TRPM7 in both adaptive and innate immunity (Mendu et al., 2020). TRPM7 plays a salient role in macrophage activation in response to inflammatory stimuli and accordingly, when *Trpm7* is deleted selectively in myeloid cells, the mice are highly resistant to endotoxemia (Schappe et al., 2018). *Trpm7^−/−^* macrophages are also defective in efferocytosis (Schappe et al., 2022), a process through which cell corpses are cleared, but they are normal in their ability to clear bacteria. Since microglia, the brain-resident macrophages, are crucial mediators of neuroinflammation, we reasoned that small molecule inhibitors of TRPM7 that accumulate in the brain may be of value in arresting neuroinflammation. Advancing this approach requires the identification of generally safe TRPM7 inhibitors that infiltrate the CNS.

TRPM7 assembles as homotetramers, that are inhibited by [Mg^2+^]_i_, and can be observed electrophysiologically as a steep, outward-rectifying, cationic current on the plasma membrane (Duan et al., 2018, Nadezhdin et al., 2023). TRPM6 is a closely related channel that can form heterotetramers with TRPM7 (Chubanov et al., 2004), decreasing its sensitivity to physiological levels of [Mg^2+^]_i_ (Ferioli et al., 2017). In contrast to the ubiquity of TRPM7, TRPM6 is primarily expressed in the kidneys and intestines (Groenestege et al., 2006) and nearly undetectable in macrophages (Heng et al., 2008). Consistently, deletion of *Trpm7* alone is sufficient to silence the outwardly rectifying, Mg^2+^-inhibitable current typical of I_M7_/I_M6_ in macrophages (Schappe et al., 2022). In addition to its presence on the plasma membrane (PM), the bulk of the TRPM7 protein is found in intracellular vesicles termed M7-vesicles or simply, M7Vs (Abiria et al., 2017). The dynamics of TRPM7 membrane trafficking in macrophages has not been studied yet and the precise contributions of PM-resident and vesicular TRPM7 in macrophage functions are still a topic of investigation.

There are a number of reported TRPM7 inhibitors, but many lack specificity or potency, being repurposed drugs with greater activity at their primary targets (Chubanov et al., 2014, Chubanov et al., 2017). NS8593 and FTY720 (Fingolimod) are such examples. NS8593 was originally developed as an inhibitor of small-conductance Ca^2+^-activated K+ (SK) channels (Strøbæk et al., 2006), but was later found to inhibit TRPM7 with an IC_50_ of 1.6 µM (Chubanov et al., 2012). However, NS8593 acts on SK-family channels with an IC_50_ in the range of 0.42 – 0.73 µM and has been shown to prolong the atrial action potential (Diness et al., 2010), limiting *in vivo* use. Waixenicin A, which is a natural compound isolated from marine soft coral has been reported to inhibit TRPM7. This study claimed an IC_50_ of 16 nM, but this IC_50_ was derived in recording conditions that limit the development of TRPM7 currents. I_TRPM7_ is best resolved in Mg^2+^-free conditions and in such conditions, the IC_50_ was shown to be 7 μM (Zierler et al., 2011). Moreover, inhibition of TRPM7 by 10 μM Waixenicin is very slow, taking up to 200 seconds to suppress I_TRPM7_. Although Waixenicin has been used *in vivo* (Turlova et al., 2021, Sun et al., 2020), there are no pharmacokinetic studies proving that it is bioavailable in blood and tissues at concentrations needed to inhibit TRPM7. Until recently, this compound could only be derived from its natural sources, and was not commercially available. However there is now a viable total chemical synthesis (Steinborn et al., 2023) and this will facilitate further studies to clarify its utility *in vivo*. Despite these advances, there remains a need for chemically accessible TRPM7 inhibitors with pharmacological properties fit for targeting TRPM7 in pre-clinical *in vivo* studies. The present study focuses on improving FTY720 and molecules related to it, as they offer the key advantages of oral administration, low toxicity, and tissue bioavailability at concentrations that can inhibit TRPM7 effectively.

FTY720 is an immunosuppressive drug that is a structural analog of sphingosine, and is currently used for the treatment of relapsing-remitting multiple sclerosis (Brinkmann et al., 2010). FTY720 is a prodrug that is phosphorylated *in vivo* by sphingosine kinases (SPHK1 and SPHK2) to yield the active immunosuppressant, FTY720-phosphate (FTY720-P) (Paugh et al., 2003). FTY720-P is a picomolar (pM) agonist of multiple Sphingosine-1-Phosphate Receptors (S1PRs), most prominently S1P1. Since supraphysiological agonism results in effective downregulation of S1P1, FTY720 acts as a functional antagonist. *In vivo*, disruption of S1P1 signaling leads to lymph node sequestration of lymphocytes and concomitant lymphopenia (Brinkmann et al., 2002, Chiba et al., 1998, Matloubian et al., 2004). Interestingly, the non-phosphorylated form of FTY720 and sphingosine have no effect on S1PRs, but both were found to inhibit TRPM7 with IC_50_s of 0.72 µM and 0.59 µM, respectively (Qin et al., 2013). Conversely, the phosphorylated form, FTY720-P, has no effect on I_M7_. Prior pharmacokinetic studies of FTY720 have demonstrated that the compound reaches tissue concentrations much higher than those observed in the blood, with lung tissue displaying the highest partition coefficient of all examined organs (Meno-Tetang et al., 2006). FTY720 has also been shown to cross the blood brain barrier and accumulate in white matter at levels between 10- and 27-times that of the blood (Foster et al., 2007). We reasoned that non-phosphorylatable analogs of FTY720 could be candidates for more selectively inhibiting TRPM7 at the exclusion of S1PR targeting; and their likely tissue distribution would make them especially well suited to *in vivo* neuropharmacology.

Although clinically efficacious, all S1P1 receptor agonists (four are marketed currently) cause initial dose bradycardia in humans, a significant deleterious side-effect that emerges from on-target S1P1 agonism (Camm et al., 2014). This species-specific effect was originally shown to be mediated by S1P3 agonism in mice (Sanna et al., 2004), which had prompted a search for S1P1-selective FTY720 analogs. These efforts led to the discovery of molecules with conformationally constrained headgroups and pre-existing chirality at their phosphorylation sites (Fig. S1) (Lynch and Macdonald, 2006). While these medicinal chemistry efforts did not yield intended drug candidates (Albert et al., 2005), they yielded a set of well-characterized small molecules that proved useful for our study. We turned our attention to AAL-149 and VPC01091.4 which are non-phosphorylatable FTY720 analogs (Fig. 1) that are not active at S1P receptors, and, when administered to mice, do not evoke lymphopenia (Brinkmann et al., 2002, Zhu et al., 2007). In the process, we have identified VPC01091.4 as a potent inhibitor of TRPM7 ion channel activity. VPC01091.4 accumulates in the brain and lungs without evidence of toxicity and in a mouse model of endotoxemia, it blunts systemic peripheral inflammation as well as neuroinflammation. An important caveat to these findings is that the anti-inflammatory effects of TRPM7 inhibitors (VCP4, AAL-149, FTY720 and NS8593) do not entirely depend on TRPM7, though the additional contributing off-target effectors are unknown. Nonetheless, VPC01091.4 represents a significant improvement in selectivity over FTY720, as it allows attempts at *in vivo* targeting of TRPM7 without driving lymphopenia, a typical result of SIPR targeting. Future work may identify more potent and selective TRPM7 inhibitors within the space of non-phosphorylatable sphingosine isomers.

**Figure 1:**
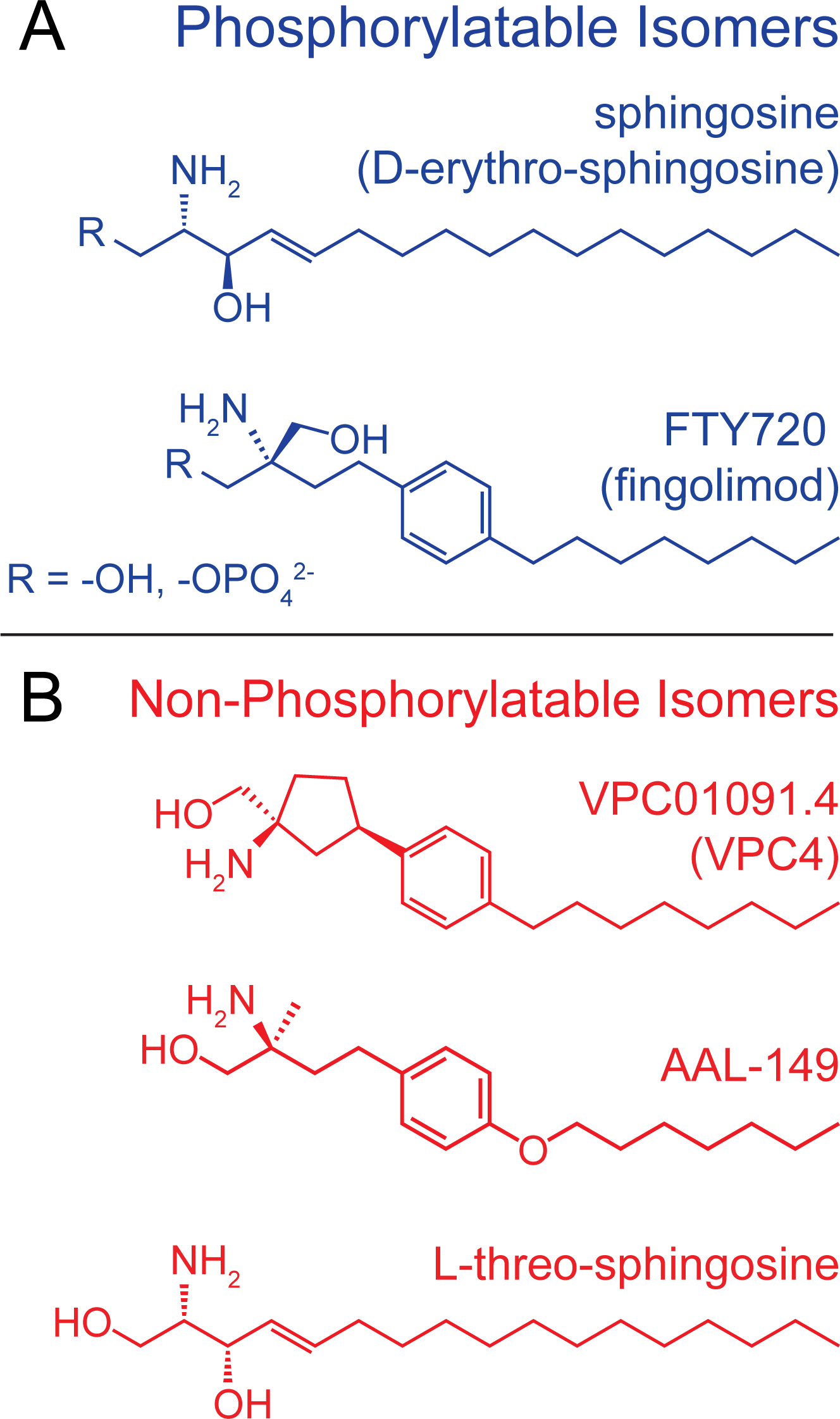
Structure of sphingosine analogs that inhibit I_TRPM7_. (A) The structure of the predominant, naturally occurring sphingosine isomer, (D-erythro-sphingosine or “sphingosine”) and FTY720 (fingolimod) are shown. Both sphingosine and FTY720 inhibit I_M7_ but they are phosphorylated *in vivo,* and the phospho-analogs are potent agonists of the ubiquitous S1P1 receptor. FTY720 is a prochiral molecule, only the *S* enantiomer (shown) is produced by sphingosine kinases. Phosphorylatable isomers are shown in blue. (B) VPC01091.4, AAL-149, and L-threo-sphingosine are structural analogs of sphingosine that are not phosphorylated *in vivo* and lack activity against S1PRs. The existing stereochemistry of these compounds places their hydroxymethyl groups in a position that is not amenable to phosphorylation by sphingosine kinases.

## Results

### Identification of VPC01091 stereoisomers that block I_M7_

VPC01091 has four stereoisomers (denoted VPC01091.1 – 4). Isomers 2 and 4 were of primary interest to us because they are not substrates of sphingosine kinases (Fig. S1B). We carried out whole-cell patch clamp electrophysiology on HEK 293T cells overexpressing mouse TRPM7 and isolated the typical outwardly rectifying I_M7_ reversing at 0 mV. I_M7_ is inhibited by free Mg^2+^ and consequently, I_M7_ is revealed using a pipette solution rich in divalent chelators that deplete free [Mg^2+^]_i_. When applied to maximal I_M7_, VPC01091.4 resulted in significant inhibition (Fig. 2A, top). VPC01091.1 and VPC01091.3 also inhibited I_M7_ (Fig. 2D & F). VPC01091.2 was not available for testing, but predictably, the application of its chemically phosphorylated variant, VPC01091.2-P, had negligible effect on peak I_M7_ (Fig. 2E). Next, by carrying out dose-response studies, we determined the IC_50_ of VPC01091.4 to be 0.665 µM for TRPM7 inhibition (Fig. 2A, bottom). Henceforth, we refer to VPC01091.4 as simply VPC4.

**Figure 2:**
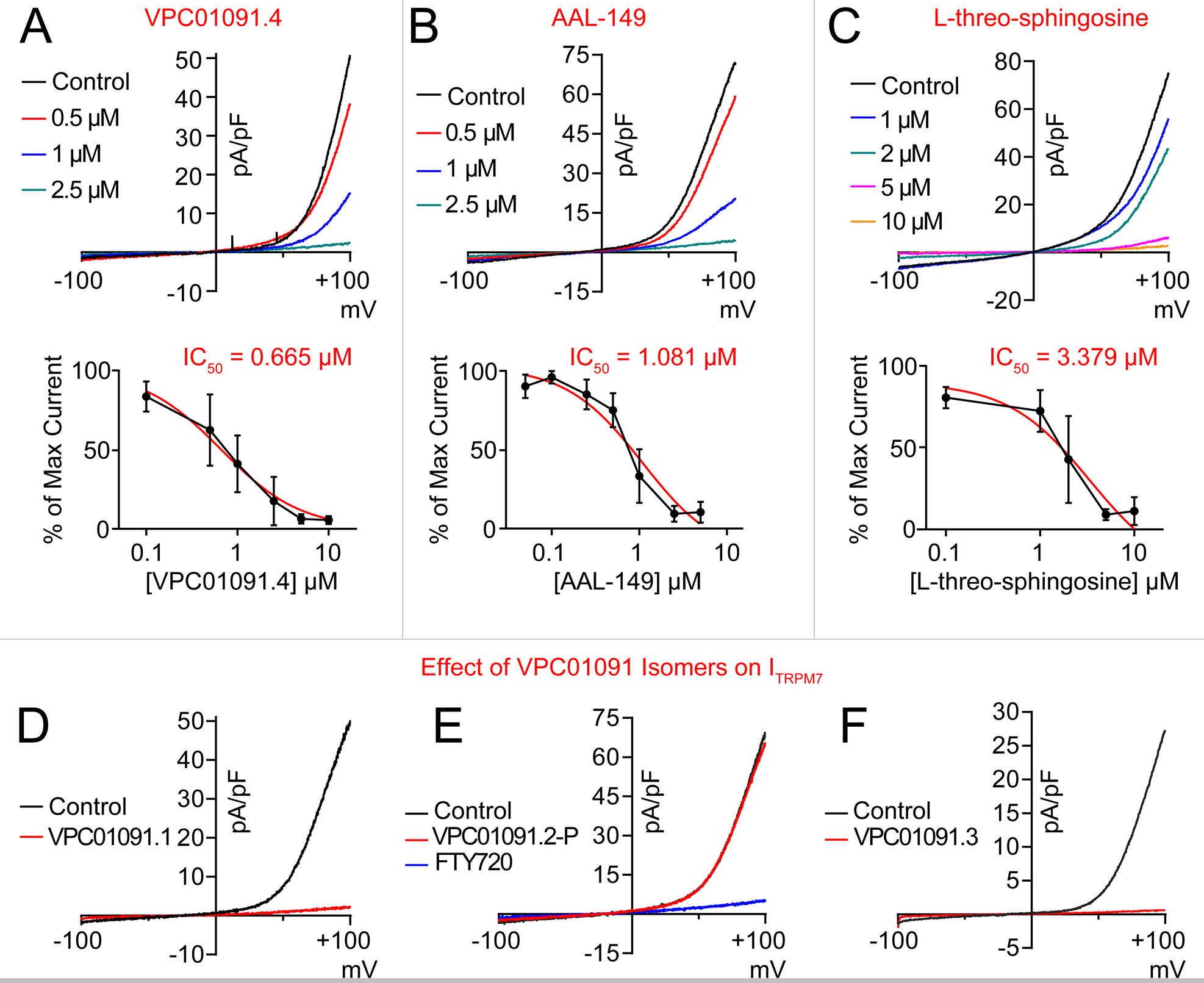
VPC01091.4, AAL-149, and L-threo-sphingosine inhibit TRPM7 current. The effects of sphingosine analogs on whole-cell TRPM7 currents from HEK 293T cells overexpressing mouse TRPM7 are shown. Peak current traces (black) represent the maximal current observed at +100 mV after allowing for I_M7_ to run-up in control bath solution and drug-treated traces represent the minimal current observed after drug application. (A – C) Representative recordings show the dose-dependent inhibition of I_M7_ by VPC01091.4, AAL-149, and L-threo-sphingosine. (A) (Top) Dose-dependent inhibition by VPC01091.4 of maximal TRPM7 currents. (Bottom) IC_50_ was determined to be 0.665 µM by best fit non-linear regression model (red trace) (95% CI IC_50_ = 0.525 to 0.834, n = 8 for each condition, SEM are shown). (B) (Top) Dose-dependent inhibition of I_M7_ by AAL-149. (Bottom) IC_50_ was determined to be 1.081 µM (95% CI IC_50_ = 0.656 to 1.866, n = 3 at 0.1 µM and n => 5 for all other conditions, SEM are shown). (C) (Top) Dose-dependent inhibition of I_M7_ by L-threo-sphingosine. (Bottom) IC_50_ was determined to be 3.379 µM (95% CI IC_50_ = 1.169 to 14.940, n = 2 at 10 µM and n => 3 for all other conditions, SEM are shown). (D – F) The effects of VPC01091 stereoisomers on whole-cell TRPM7 currents from HEK 293T cells overexpressing mTRPM7 are shown. Example traces from cells treated with 10 µM VPC01091.1 (D) or VPC01091.3 (F). VPC01091.2 was not available to be tested, but the chemically phosphorylated VPC01091.2-P had no effect on IM7 (E), consistent with the observation that S1P and FTY720-P do not inhibit I_M7_. Subsequent application of 5 µM FTY720 was able to inhibit I_M7_.

### AAL-149 and L-threo-sphingosine are additional blockers of I_TRPM7_

Next, we evaluated an additional sphingosine-like compound, AAL-149, which is the non-phosphorylatable enantiomer of AAL-151 (Fig. S1C). AAL-149 also inhibited I_M7_ in a dose-dependent manner, with a calculated IC_50_ of 1.081 µM (Fig 2B). VPC4 and AAL-149 thus represent drug candidates with similar potency against TRPM7 as FTY720 (IC_50_ = 0.72 µM). However, these proprietary compounds are not commercially available. For that reason, we also attempted to identify a more widely available analog of sphingosine that cannot be phosphorylated. “Sphingosine,” or D-erythro-sphingosine (*2S,3R*), is one of 4 possible stereoisomers, and the predominant naturally occurring isomer (Merrill, 2002). We took advantage of the observation that while the erythro-enantiomers are phosphorylated by mammalian sphingosine kinases, the threo-enantiomers are competitive inhibitors of sphingosine kinases and not appreciably phosphorylated *in vivo (Buehrer and Bell, 1992)*. We tested a commercially available L-threo-(*2S,3S*)-sphingosine for its potential to inhibit I_M7_ (Fig. 1B). We found that L-threo-sphingosine inhibits TRPM7 current (Fig. 2C), with a reduced potency in comparison to VPC4, AAL-149, and FTY720. The calculated IC_50_ was 3.379 µM. For subsequent cell biological studies, we focused on VPC4 and AAL-149.

### VPC4 and AAL-149 are not cytotoxic at concentrations necessary to inhibit I_M7_

To test whether VPC4 and AAL-149 could be tolerated at effective inhibitory doses, we incubated HeLa cells for 24 hours with a range of compound concentrations and measured LDH release as a readout of cytotoxicity (Fig. 3A). For both VPC4 and AAL-149, dosages up to 10 µM did not result in any significant cytotoxicity. At the highest dose of 25 μM, which is more than 30X its IC_50_, VPC4 treated cells exhibited modest cytotoxicity (4% compared to positive control of 100% cytotoxicity). These findings establish VPC4 and AAL-149 as viable alternatives to FTY-720 for cell-based investigations of TRPM7, with the key advantage that they do not act on S1P receptors (Brinkmann et al., 2002, Zhu et al., 2007).

**Figure 3:**
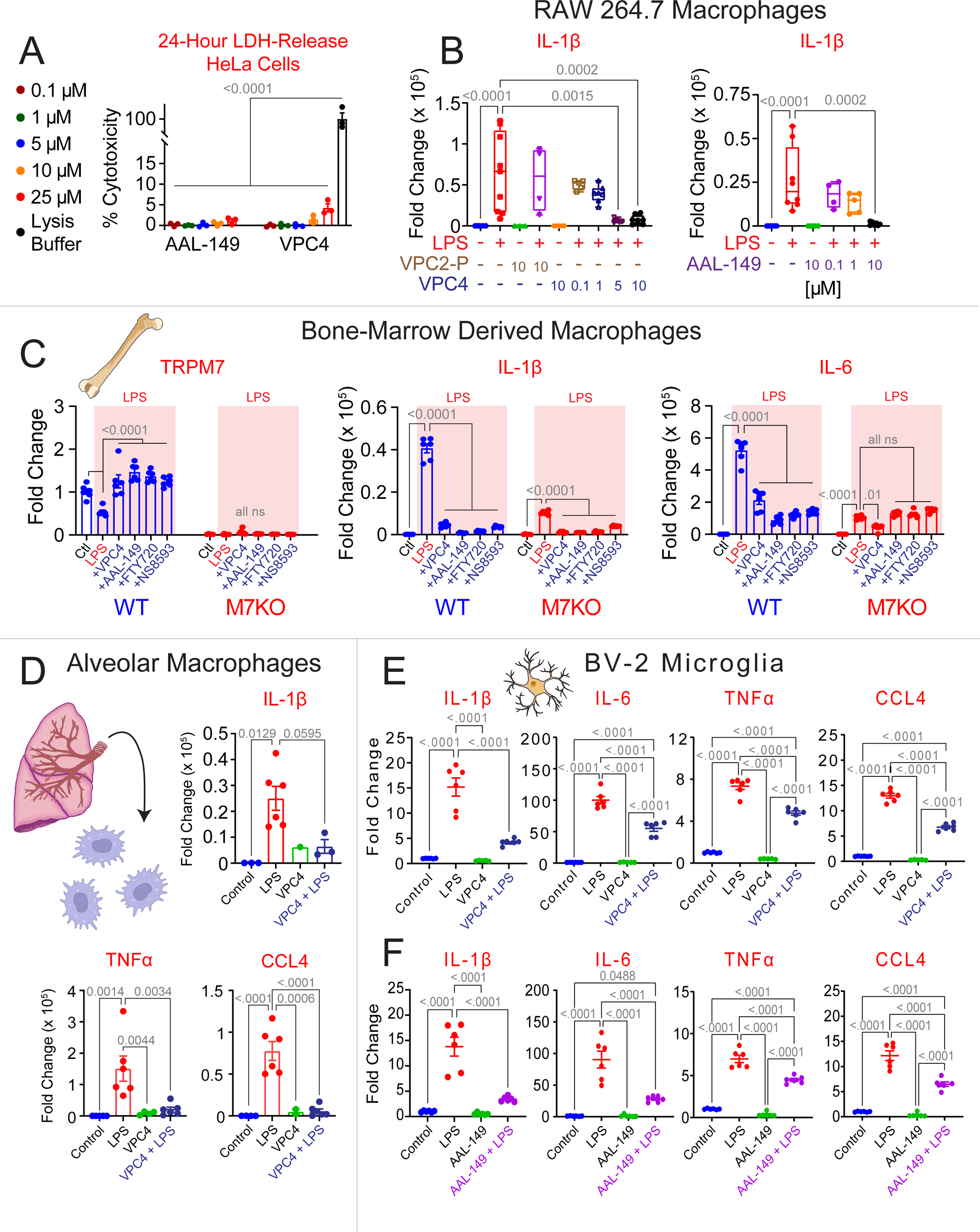
VPC01091.4 and AAL-149 suppress LPS-induced inflammation in macrophages and microglia. (A) HeLa cells display minimal cytotoxicity to AAL-149 and VPC01091.4 at doses up to 25 µM (mean cytotoxicity <5% for each condition, n = 3 for each dose). P< 0.0001 for all comparisons relative to lysis control. (B) Change in *IL-1β* expression observed by qRT-PCR in RAW 264.7 macrophages following stimulation with LPS (1 µg/mL for 3 hours). Pretreatment with >=5 µM of VPC01091.4 or 10 µM AAL-149, but not VPC01091.2-P, significantly reduced LPS-stimulated *IL-1β* expression. For brevity, only statistically significant comparisons from the LPS-only treated groups are shown. Box and Whisker plots display the minimum, first quartile, median, third quartile, and maximum values. (C) Bone-Marrow Derived Macrophages (BMDMs) from wildtype (*Trpm7^fl/fl^*) and TRPM7 KO (*Trpm7^fl/fl^ LysM Cre*) mice were isolated and differentiated for 7 days prior to use for an LPS-induced inflammation assay. Expression levels of *Trpm7*, *IL-1β*, and *IL-6* as measured by qRT-PCR are displayed. Levels of TRPM7 expression are normalized to control-treated WT BMDMs, and *IL-1β*, and *IL-6* expression levels are normalized to control-treated BMDMs for their respective genotype. 4 different TRPM7 inhibitors were used: 10 µM VPC4, 10 µM AAL-149, 5 µM FTY720, or 30 µM NS8593. Statistical analysis was performed using a 2-way ANOVA with Tukey’s multiple comparison within each genotype. (D) Alveolar macrophages isolated from wildtype mice were stimulated with LPS as above. Pre-treatment with 10 µM VPC4 significantly suppressed expression of *IL-1β*, *TNFα*, and *CCL4*. (E & F) Inflammatory cytokine expression levels as measured by qRT-PCR in cultured BV-2 microglia pre-treated with vehicle, 10 µM VPC01091.4 (E), or 10 µM AAL-149 (F) before stimulation with LPS (1 µg/mL for 3 hours). Unless stated otherwise, ordinary one-way ANOVA were used for analyses and error bars represent SEM. The vehicle used for VPC4, VPC2-P, FTY720, and NS8593 was DMSO and the vehicle for AAL-149 was 100% ethyl alcohol (<0.2% v/v in experimental media for each). VPC4 = VPC01091.4 and VPC2-P = VPC01091.2-P.

### VPC4 and AAL-149 inhibit LPS-induced inflammatory gene expression in macrophages and microglia

We have previously shown that *Trpm7^−/−^* macrophages exhibit significantly reduced inflammatory gene expression in response to LPS and other TLR ligands (Schappe et al., 2018). Based on the use of FTY720, we also concluded that the ion channel activity of TRPM7 was paramount for its function in inflammatory signaling but acknowledged the caveat that at the concentrations used, FTY720 would clearly target the macrophage S1PRs. We therefore tested whether VPC4 and AAL-149 can inhibit macrophage activation in response to LPS. RAW 264.7 macrophages were pretreated with either VPC4, VPC01091.2-P, or AAL-149, prior to exposure to LPS (1 µg/mL for 3 hours) (Fig. 3B). Macrophages stimulated with LPS displayed a significant upregulation in *IL-1β* expression relative to unstimulated macrophages. When pre-treated with VPC01091.2-P, which does not inhibit TRPM7, there was no significant inhibition of *Il-1β* gene expression. However, in cells pretreated with VPC4, the gene expression of *Il-1β* was almost completely abrogated at 5 µM and 10 µM. Similarly, pre-treatment with AAL-149 (10 µM) also completely abrogated *Il-1β* gene expression in response to LPS.

Next, we sought to evaluate the TRPM7-dependent anti-inflammatory effects of VPC4 and AAL-149 and compare them to the widely used TRPM7 inhibitors FTY720 and NS8593. (Fig. 3C). Bone-Marrow Derived Macrophages (BMDMs) from WT (*Trpm7^fl/fl^*) and TRPM7 KO (*Trpm7^fl/fl^ LysM Cre*) mice were isolated and differentiated for 7 days prior to use for an LPS-induced inflammation assay. TRPM7 KO BMDMs had near total loss of *Trpm7* expression relative to WT cells (Fig. 3C, left). Interestingly, WT BMDMs demonstrated a significant reduction in *Trpm7* expression after treatment with LPS, and this effect was prevented with treatment of any of the 4 tested TRPM7 inhibitors. Next, we observed that LPS-stimulated expression of *Il-1β* and *IL-6* was greatly reduced in the TRPM7 KO BMDMs relative to WT. In *Trpm7* KO BMDMs, *IL-6* expression was suppressed to similar degrees both genetically and pharmacologically, with the exception of VPC4-treatment where an additive effect was observed (Fig. 3C, right). In both, TRPM7 WT and KO BMDMs, *Il-1β* expression was suppressed through treatment with the TRPM7 inhibitors (Fig. 3C, middle). Taken together, these data indicate that the majority of LPS-induced *Il-1β* expression is driven in a TRPM7-dependent manner, though the remainder is sensitive to the pharmacologic inhibitors, suggestive of an additive off-target effect of these drugs.

Lastly, we observed that VPC4 exhibited direct anti-inflammatory effects upon primary mouse alveolar macrophages (Fig. 3D) and the mouse BV-2 microglial cell line (Fig. 3E). AAL-149 also exerted similar anti-inflammatory effects (Fig. 3F). These results recapitulate the findings from reverse genetic studies and establish VPC4 and AAL-149 as small molecule inhibitors of TRPM7 ion channel activity with potent anti-inflammatory effects. The findings also bolster the critical role of TRPM7 ion channel activity as a major driver of LPS-driven macrophage inflammatory signaling.

### VPC4 reduces systemic peripheral inflammation in a mouse model of LPS-induced endotoxemia

Previously, we showed that a myeloid-specific knockout of *Trpm7* (*Trpm7^fl/fl^ LysM Cre*) reduced systemic peripheral inflammation in a mouse model of LPS-induced peritonitis (Schappe et al., 2018). However, the potential of TRPM7 as an immunomodulation target has not been validated through pharmacological approaches. Additionally, since TRPM7 is required for embryogenesis and expressed widely, there have been lingering doubts that drugging TRPM7 would result in systemic toxicity, negating its potential in immunomodulatory therapies. The identification and characterization of VPC4 as a TRPM7 inhibitor that does not result in significant cytotoxicity allowed us to test whether pharmacologically targeting TRPM7 would suppress endotoxin mediated inflammation without undue systemic toxicity. We used a sub-lethal, LPS-induced peritonitis model for systemic endotoxemia in mice. 12-week-old, wildtype mice received an intraperitoneal injection with either vehicle, LPS (1 mg/kg), VPC4 (30 mg/kg) or a co-injection of VPC4 and LPS. They were monitored for 4 hours for overt pathological symptoms. After 4 hours, the mice were euthanized for collection of whole blood and tissues (Fig. 4A). All mice which received VPC4 achieved whole blood levels that were well above the IC_50_ for TRPM7 blockade (0.665 µM), with mean concentrations of 15.78 µM and 17.35 µM for the drug-treated and co-treatment groups, respectively (Fig. 4B). Using an automated hematology analyzer, we measured absolute lymphocyte counts and observed no significant differences between the treatment groups, in line with a previous demonstration that VPC4 does not drive lymphopenia in mice (Zhu et al., 2007) (Fig. 4C). Luminex analysis of plasma cytokines revealed a significant elevation in the levels of inflammatory cytokines *IL-1β*, *IFNγ*, and *TNFα* in LPS-treated mice. These inflammatory cytokines were significantly suppressed in mice co-treated with LPS and VPC4 (Fig. 4D). The co-treatment group also demonstrated higher levels of the anti-inflammatory cytokine *IL-10*.

**Figure 4:**
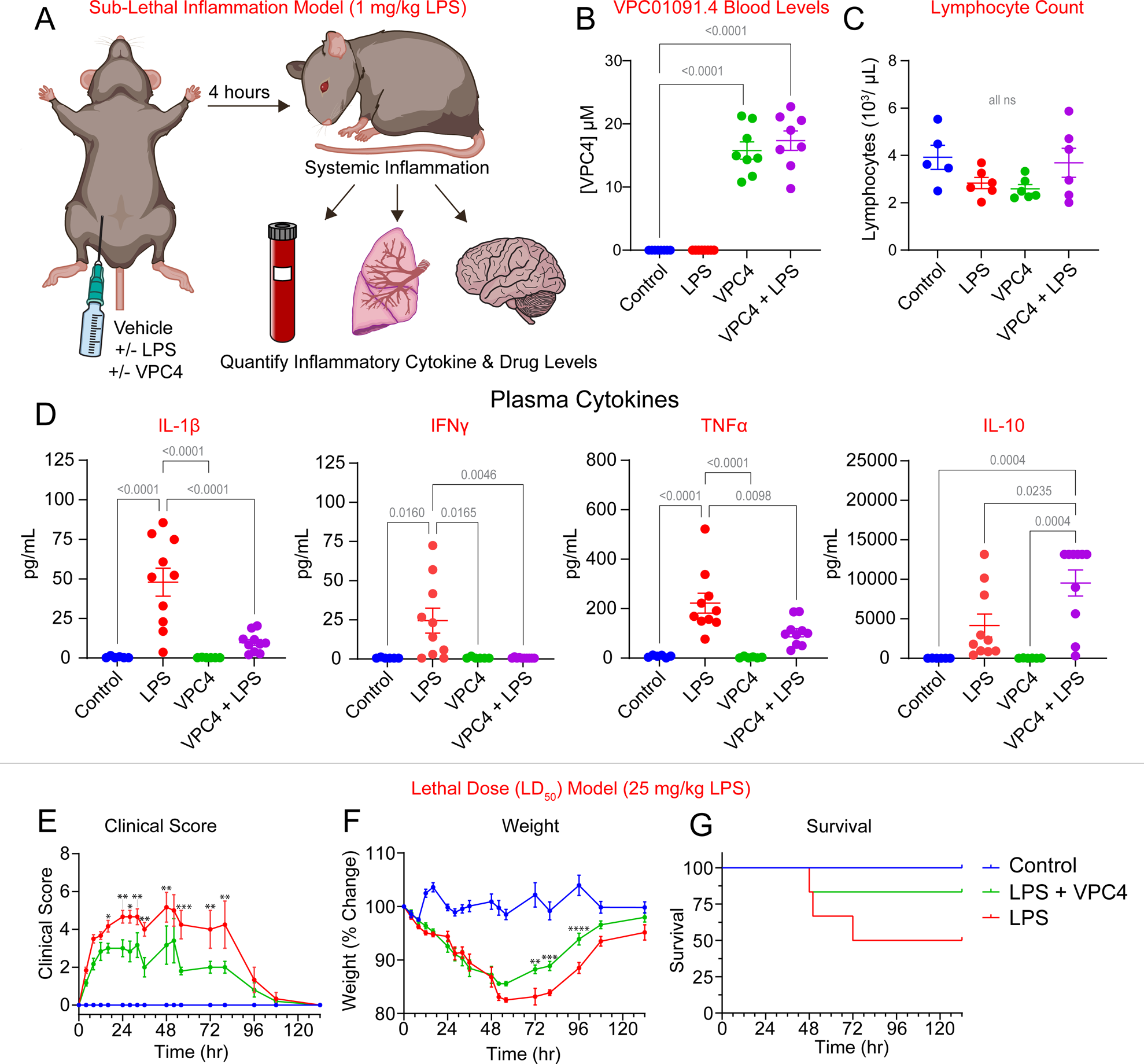
VPC01091.4 suppresses the LPS-induced systemic inflammatory response *in vivo*. (A) Experiment Schematic for the sub-lethal, LPS-induced inflammation model: 12-week-old, wildtype C57BL/6 mice were injected intraperitoneally with LPS (1 mg/kg) with or without co-administered VPC01091.4 (30 mg/kg). After 4 hours, mice were euthanized and blood was collected for determination of whole blood drug levels, lymphocyte counts, and plasma cytokine levels. Following biventricular perfusion, lung and brain tissue were collected (see organ data in Figure 5). (B) Whole-blood levels of VPC01091.4 as determined by LC-MS. Ordinary one-way ANOVA tests were performed for statistical analysis, SEM are shown. (C) Mean lymphocyte counts as determined by an automated hematology analyzer. Ordinary one-way ANOVA tests were performed for statistical analysis, SEM are shown. (D) Plasma cytokines as determined by a Luminex assay for *IL-1β*, *IFNy*, *TNFα*, and *IL-10*. Data are pooled from 4 independently conducted trials with 8 mice each (n = 6 for Control and VPC4-only groups and n = 10 for LPS and VPC4 + LPS groups). Ordinary one-way ANOVA tests were performed for statistical analysis, SEM are shown. (E – G) Data from a lethal dose (LD_50_) LPS Model. The experiment was performed similarly to the sub-lethal model, except mice received an LD_50_ dose of LPS (25 mg/mL), with or without co-administered VPC01091.4 (30 mg/kg); then monitored for 132 hours. Weights and clinical symptoms were measured at each timepoint. (E) Clinical scores over time by treatment group. Significant differences between the LPS and LPS + VPC4 treated groups as determined by a Mixed-Effects ANOVA analysis are shown (* = p ≤ 0.05, ** = p ≤ 0.01, *** = p ≤ 0.001). (F) Changes in weight from baseline over time by treatment group. Significant differences between the LPS and LPS + VPC4 treated groups as determined by a Mixed-Effects ANOVA analysis are shown (** = p ≤ 0.01, *** = p ≤ 0.001, **** = p ≤ 0.0001). (G) Kaplan-Meier survival curve by treatment group. The survival rate was 100% for the control group, 50% for the LPS-treated group, and 83.33% for the LPS + VPC4 co-treated group (n = 6 mice for each group). A log-rank Mantel-Cox test determined there were no significant differences between treatment groups, with a p-value of 0.127.

To test whether VPC4 treatment could increase the survival of mice in the setting of LPS-induced endotoxemia, we challenged mice with a 50% lethal dose of LPS and monitored their weight and symptoms for 132 hours. The severity of observed conjunctivitis, lethargy, grooming quality, stool changes, and facial grimace were assessed based on standardized scoring parameters (Table 1) to obtain composite clinical scores (Fig. 4E). Mice which receive LPS treatment exclusively had the most severe symptoms, and these symptoms were significantly reduced at a number of timepoints in mice which were co-treated with LPS and VPC4. Control mice displayed no adverse effects at any time points after receiving vehicle treatment. LPS-only and co-treated mice lost weight to a similar degree through the first 48 hours of the experiment, at which point the co-treated mice began recovering at a higher nadir and at an earlier timepoint as compared to the LPS-only treated mice (Fig. 4F). The survival rate of LPS-treated mice was 50%, compared to a survival rate of 83.33% in the co-treated mice and 100% in the control mice (Fig. 4G); though the differences were not statistically significant with a p-value of 0.127. At the final timepoint, all mice which had survived the experiment had returned to baseline symptoms with no remaining evidence of toxicity.

**Table 1:** Scoring Sheet for Humane Endpoints: LPS-Induced Peritonitis Model.

| Scoring Parameter: |  | 0 | 1 | 2 |
| --- | --- | --- | --- | --- |
|  | Conjunctivitis | Normal | Single eye open with visible discharge | Eyes closed with discharge and swelling |
|  | Lethargy | Normal locomotion and reaction, >3steps | Inactive, <3 steps after moderate stimulation, slight hunching<br>**SCORE AS 2 FOR THIS CATEGORY** | Only lifting of head after moderate stimulation <1 step, severe hunching<br>**SCORE AS 4 FOR THIS CATEGORY** |
|  | Hair Coat | Well-groomed with smooth coat | Rough coat, minor ruffling | Unkempt fur, dull coat |
|  | Grimace Pain | Normal | Moderate orbital tightening or nose bulge | Sever orbital tightening, nose bulge, and collapsed ear position |
|  | Stool | Normal | Creamy consistency | Watery diarrhea |
| Animals receiving a **total score equal to or greater than 8** or those which have **lost 20% of more of their body weight** will be euthanized. |  |  |  |  |

Taken together, VPC4 appeared to be well-tolerated and capable of suppressing the systemic inflammatory response produced by intraperitoneal LPS injection without driving lymphopenia. In addition, it reduced the severity of physical symptoms and weight loss in mice challenged with a lethal dose of LPS and produced a non-significant increase in mouse survival.

### VPC4 accumulates in the brain and lungs

FTY720 broadly distributes throughout the body, including the central nervous system, and reaches the highest concentrations in the lungs (Meno-Tetang et al., 2006). A later study showed more specifically that FTY720 passes through the BBB and accumulates in the white matter (Foster et al., 2007). We were hopeful that VPC4 would exhibit similar pharmacological properties and thus prove useful for modulating brain inflammation. Lung and brain tissues collected during our LPS-induced peritonitis model were partitioned, with a portion of the tissue preserved immediately for whole-organ RNA extraction. The remaining tissue was weighed, then mechanically homogenized in PBS with an internal standard (0.5 µM FTY720). The homogenate was then analyzed by LC-MS. Based on the LC-MS signal intensities of VPC4 and the internal standard (FTY720), the VPC4 concentrations were calculated. In the mice that received VPC4, the VPC4 levels in the brain were approximately 9-fold higher than those found in the blood (Fig. 5A). Lung levels of VPC4 were even higher, reaching 32-fold higher than the blood levels (Fig. 5C). These results confirm that, like FTY720, VPC4 can pass through the BBB. We then evaluated whether VPC4 dampened inflammation in the lung and brain.

**Figure 5:**
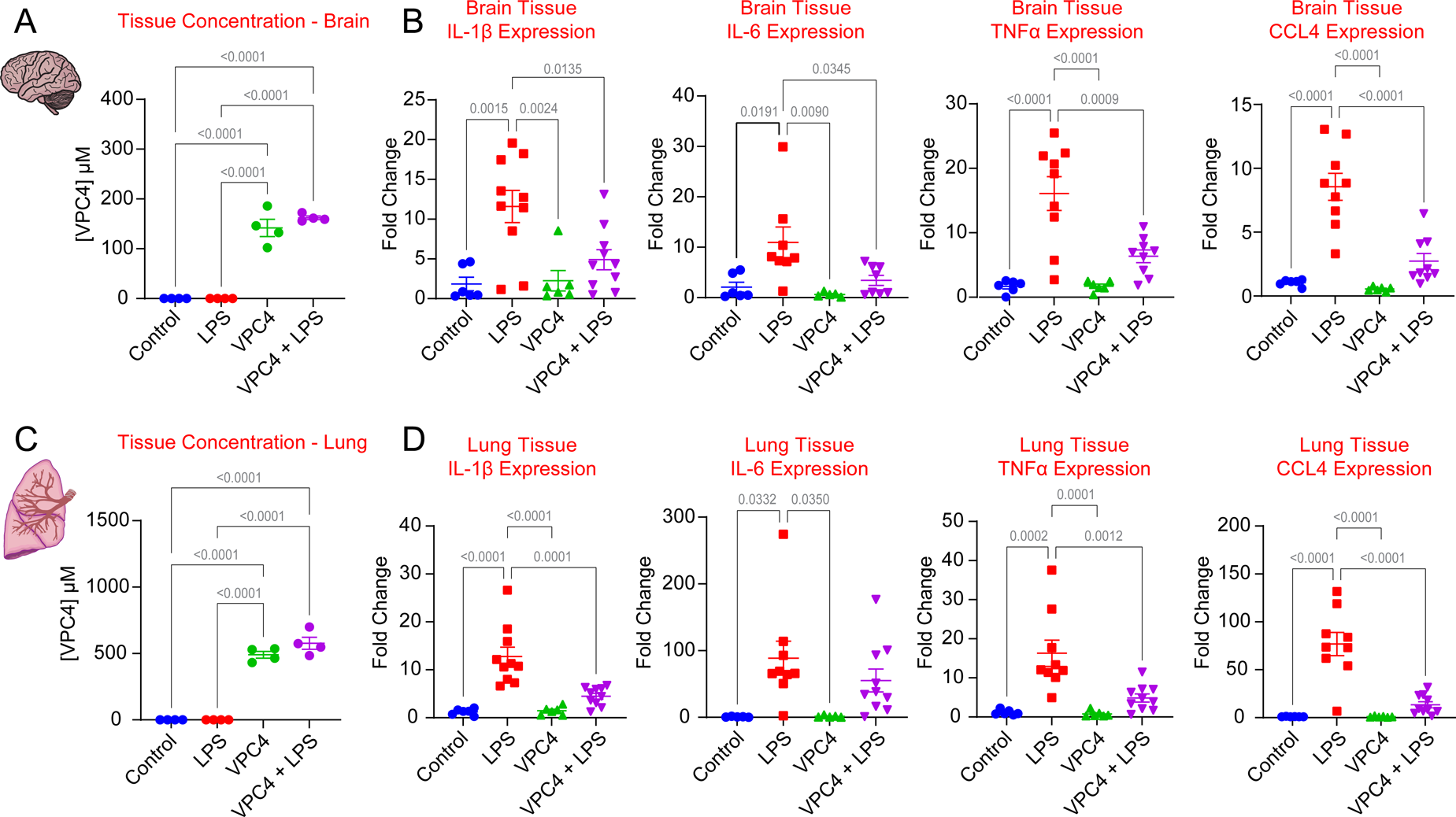
VPC01091.4 crosses the blood brain barrier and exerts organ-level anti-inflammation in lung and brain tissue. (A) Whole-brain levels of VPC01091.4 as determined by LC-MS. Drug-treated groups had mean brain concentrations of 141.8 µM (VPC4) and 162.2 µM (VPC4 + LPS), 8.97- and 9.35-times higher than mean blood levels, respectively. (B) Expression of inflammatory cytokines as measured by qRT-PCR of RNA extracted from whole-brain homogenates. (C) Whole-lung levels of VPC01091.4 as determined by LC-MS. Drug-treated groups had mean lung concentrations of 490.3 µM (VPC4) and 576.3 µM (VPC4 + LPS), 31.07- and 33.22-times higher than mean blood levels, respectively. (D) Expression of inflammatory cytokines as measured by qRT-PCR of RNA extracted from whole-lung homogenates. Ordinary one-way ANOVA tests were performed for statistical analysis, SEM are shown.

### VPC4 blunts brain and lung inflammation in the mouse model of endotoxemia

It is well established that peripherally administered endotoxin rapidly induces inflammatory cytokines in the brain (Pitossi et al., 1997, Layé et al., 1994, Gatti and Bartfai, 1993). Whole-tissue RNA was extracted from lung and brain samples and analyzed using qRT-PCR for differences in inflammatory cytokine expression between the treatment groups. Mice treated with LPS had significantly higher brain expression of *IL-1β*, *IL-6*, *TNFα*, and *CCL4* as compared to control mice, and these cytokines were significantly suppressed through co-treatment with VPC4 (Fig. 5B). A similar effect was observed in the lung tissue, except for *IL-6*, where the suppression by cotreatment with VPC4 was non-significantly lower than the LPS group (Fig. 5D). Overall, these findings establish VPC4 as a small molecule inhibitor of TRPM7 that can pass the blood-brain barrier and blunt whole-brain and lung inflammation during systemic LPS-driven inflammation.

## Discussion

Because of their easy accessibility on the cell membrane and rapid switch-like activity that can be locked by small molecules and peptides into ON or OFF states, ion channels are the preferred molecular targets of venoms in nature and many clinical drugs (Wulff et al., 2019). Ion channels expressed by the immune cells are therefore emerging as important targets for immunomodulation (Feske et al., 2015, Froghi et al., 2021). Our previous work has established TRPM7 ion channel activity as a regulator of macrophage-mediated inflammation but a lack of effective small molecule inhibitors that are well-tolerated *in vivo* prevented pharmacological exploration. This study took advantage of the earlier finding that FTY720, a FDA-approved drug that targets S1P1 for the treatment of multiple sclerosis, inhibits TRPM7 in its unphosphorylated pro-drug form, but not in its phosphorylated form. The approach has identified VPC01091.4 (VPC4) as an effective inhibitor of TRPM7 that is well-tolerated in mice and has a potent anti-inflammatory effect in a mouse model of endotoxemia. Note that even at high doses, VPC4 was shown to have no deleterious effect on lymphocyte populations, in line with previous demonstration of its use *in vivo* (Zhu et al., 2007).

Microglia, the brain-resident macrophages, are major drivers of neuroinflammation and alleviating neuroinflammation may yield clinical benefit in slowing the progression of many neurodegenerative diseases (Streit et al., 2004, Muzio et al., 2021). Microglia-resident TRPM7 may therefore offer a direct pharmacological approach to blunt neuroinflammation. Our findings show that VPC4 can inhibit microglial activation *ex vivo*, pass the blood-brain barrier, and greatly blunt neuroinflammation in an endotoxemia mouse model. This study is a significant advance because it sets the stage for translational studies that test the potential of TRPM7 as a drug target in neurodegenerative diseases. Promising results in such studies may, in turn, encourage the development of a new class of more potent and selective inhibitors of TRPM7. In addition to the brain, we show that VPC4 accumulates in the lungs at especially high levels. The function of TRPM7 in lung macrophages has not been studied yet but alveolar macrophages are the most abundant immune cells in the lung (Cai et al., 2014), and significant drivers of lung immunopathologies such as acute respiratory distress syndrome (Aggarwal et al., 2014). The pharmacokinetic properties reinforced here suggests that this class of drugs may prove especially useful in targeting lung pathologies.

The mechanism through which FTY720 and VPC4 inhibit TRPM7 ion channel activity has not been explored yet. Like many TRP channels, TRPM7 ion channel activity is highly sensitive to interactions with membrane lipids. This was recently reinforced through a structural demonstration that the TRPM7 inhibitors NS8593 and VER155008 stabilize the closed state of TRPM7 through binding to a vanilloid-like pocket that likely interacts with endogenous lipids (Nadezhdin et al., 2023). Sphingosine and FTY720 are not thought to be pore blockers of TRPM7, as they reduce I_M7_ by lowering the open probability without affecting single channel conductance (Qin et al., 2013). It is therefore possible that sphingosine-like antagonists like FTY720 and VPC4 interfere with the ability of TRPM7 to interact with regulatory phospholipids (Runnels et al., 2001, Kozak et al., 2005). Future structural studies will clarify the underlying mechanisms and facilitate a rational modification of VPC4 for increased potency and selectivity. Thus, this class of non-phosphorylatable FTY720 analogs represents a fruitful search space for further optimized TRPM7 inhibitors and may lead to a new class of immunomodulatory drugs.

## Author Contributions

Conception: GWB, KRL, BND

Research Design: GWB, BND

Investigation: GWB, WHI, TH, SSN, JAK, MCM, CAD, PVS

Data analysis: GWB, WHI, TH

Resource assistance: KRL, BND

Writing - Draft and Editing: GWB, BND

Project Administration: BND

## Acknowledgements

We would like to thank all members of the Desai and Lynch labs for their scientific insights and feedback. We thank Drs. Kevin Lynch (UVA), Doug Bayliss (UVA), Stefanie Redemann (UVA), Norbert Leitinger (UVA), and Mark Beenhakker (UVA) for helpful and timely advice as Thesis Committee advisors to GWB.

The work was predominantly funded with following NIH research grants: AI155808 (BND), GM108989 (BND), R01AI144026 (KRL), Owens Family Foundation (BND), UVA MSTP Training Grant T32 GM007267 (GWB), and Pharmacological Sciences Training Grant T32 GM007055 (GWB).

The illustrations presented here are original artwork adapted from figures created using BioRender.com.

## Methods

### Resource Availability

#### Lead Contact

Further information and requests for resources and reagents should be directed to and will be fulfilled by the Lead Contact, Bimal N. Desai

#### Materials Availability

All unique/stable reagents generated in this study are available from the Lead Contact with a completed Materials Transfer Agreement.

### Cell Culture

All cells were cultured at 37°C with 5% CO_2_. HEK 293T cells (ATCC CRL-3216), HeLa cells (ATCC CCL-2), RAW 264.7 macrophages (ATCC TIB-71), and BV-2 microglial cells (AcceGen Biotech ABC-TC212S) were cultured in DMEM (Gibco 11995065) with 10% Fetal Bovine Serum (FBS, Avantor Seradigm Premium Grade Fetal Bovine Serum 97068-085). FBS was heat inactivated prior to use at 56°C for 30 minutes. Bone Marrow Derived Macrophages were isolated and cultured as previously described (Schappe et al., 2018) and differentiated into mature macrophages over 7 days through culture in BMDM media (Gibco RPMI 1640, 10% FBS, and 20% L-929 culture supernatant).

Alveolar macrophages were isolated and cultured from wildtype C57BL/6 mice (Jackson Laboratory, Strain # 000664) using an established protocol (Nayak et al., 2018). Briefly, euthanized mice were perfused with 10 mL of ice-cold PBS through the right ventricle until the lung tissue exhibited a pale-white appearance. A 22G catheter was then inserted into the trachea immediately caudal to the larynx and affixed in place using a sterile suture around the trachea. Using a 5 mL syringe filled with 3 mL of bronchoalveolar lavage buffer (“BAL buffer”: Ca^2+^- and Mg^2+^-free PBS + 1 mM EDTA), the lungs were first filled with ∼2 mL of BAL buffer, then flushed 5 times in a cycle involving the installation and removal of 1 mL of buffer. The collected BAL fluid was pooled, and the cyclical perfusion step was repeated with fresh buffer until a total of ∼10 mL of BAL fluid was collected. BAL fluid with gross evidence of blood contamination was discarded. A pellet was obtained through centrifugation at 500 x *g* for 5 minutes, contaminating red blood cells were lysed by resuspending cells for 5 minutes in 5 mL of ice-cold ACK Lysis buffer (Gibco, A1049201). The cells were then pelleted at 500 x *g* for 5 minutes, resuspended in alveolar macrophage culture media (DMEM supplemented with 10% FBS, 20% L-929 culture supernatant, 1 mM sodium pyruvate, 10 mM HEPES, and 1x penicillin/streptomycin), and plated onto untreated tissue culture dishes for 24. The media was exchanged prior to use to remove unattached cells.

To overexpress TRPM7, HEK 293T cells were plated at a density of 500,000 cells per well (day 0), transfected at 80-90% confluency (day 1), and were patched 24-36 hours post-transfection (day 2). TRPM7 overexpression was performed using wildtype mouse TRPM7 (2.5 µg plasmid DNA) (Desai et al., 2012) and jetOPTIMUS® DNA Transfection Reagent (Polyplus, 101000006). To enhance selection rate of TRPM7-overexpressing cells, eGFP (Addgene #22152) was included at a 1:10 dilution (0.25 µg) and only GFP-positive cells were patched. The observed overexpression current displayed the characteristic outward rectification, run-up, and reversal potential of I_TRPM7_ and was inhibited by 10 mM MgCl_2_ and 5 µM FTY720.

All cells were cultured at 37°C with 5% CO2. HEK 293T cells (ATCC CRL-3216), HeLa cells (ATCC CCL-2), RAW 264.7 macrophages (ATCC TIB-71), and BV-2 microglial cells (AcceGen Biotech ABC-TC212S) were cultured in DMEM (Gibco 11995065) with 10% Fetal Bovine Serum (FBS, Avantor Seradigm Premium Grade Fetal Bovine Serum 97068-085). All cell lines were negative for mycoplasma contamination as confirmed using a MycoStrip™ - Mycoplasma Detection Kit (InvivoGen rep-mys-10). FBS was heat inactivated prior to use at 56°C for 30 minutes. Bone Marrow Derived Macrophages were isolated and cultured as previously described (Schappe et al., 2018) and differentiated into mature macrophages over 7 days through culture in BMDM media (Gibco RPMI 1640, 10% FBS, and 20% L-929 culture supernatant).

### Test Compounds

**VPC01091.1** [((*1R,3S*)-1-amino-3-(4-octylphenyl)cyclopentyl)methanol] is commercially available (Avanti Polar Lipids, 857345). **VPC01091.2-P** [((*1S,3S*)-1-amino-3-(4-octylphenyl)cyclopentyl)methyl dihydrogen phosphate], **VPC01091.3** [((*1R,3R*)-1-amino-3-(4-octylphenyl)cyclopentyl)methanol], **VPC01091.4** [((*1S,3R*)-1-amino-3-(4-octylphenyl)cyclopentyl)methanol], and **AAL-149** ((2S)-2-amino-4-(4-heptoxyphenyl)-2-methylbutan-1-ol) were generously provided by Dr. Kevin Lynch. **FTY720** (fingolimod), **L-threo-sphingosine**, and **NS8593** were purchased from Cayman Chemical (Item #10006292, 10010541, and 29774, respectively). AAL-149 was solubilized in 100% ethanol, all other compounds were solubilized in DMSO.

### Patch Clamp Electrophysiology

Whole cell patch clamp electrophysiology was performed as described previously (Schappe et al., 2022). Briefly, the external solution contained (in mM): 140 Na-methanesulfonate, 5 Cs-gluconate, 2.5 CaCl_2_, 10 HEPES, pH 7.4 (adjusted with NaOH), at an osmolality of 280 – 290 mOsm/kg. The internal pipette solution contained (in mM): 115 Cs-gluconate, 3 NaCl, 0.75 CaCl_2_, 10 HEPES, 10 HEDTA, 1.8 Cs4-BAPTA, 2 Na_2_ATP, pH 7.3 (adjusted with CsOH), at an osmolality of 273 mOsm/kg. Intracellular free [Ca^2+^] was estimated to be ∼100 nM using the online WEBMAXC Standard calculator at: (https://somapp.ucdmc.ucdavis.edu/pharmacology/bers/maxchelator/webmaxc/webmaxcS.htm) (Bers et al., 2010). Recordings were captured in pCLAMP 9 (Molecular Devices) using a 400 ms ramp protocol from −100 to +100 mV and a holding potential of 0 mV. Sampling was performed at 10 kHz with a 5 kHz low-pass band filter using an Axopatch 200B amplifier (Molecular Devices). All electrophysiology experiments were conducted at RT (∼23°C). Because I_M7_ exhibits a strong flow-induced transient increase in current (Oancea et al., 2006), the maximum current is taken as the stable maximum current observed prior to the perfusion of the next solution. Similarly, the minimum current for each inhibitor dose is taken as the stable minimum current observed after the drug has had time to exert effect and after the flow-driven transient increase has subsided.

### LDH-release cytotoxicity assay

WT HeLa cells were plated at a density of 10,000 cells per well into a 96-well tissue culture microplate (Costar, 3610). AAL-149, VPC01091.4 or vehicle at 0.1, 1, 5, 10, and 25 µM was added to cells with media for a final volume of 100 µL and incubated for 24 hours. LDH release was quantified using a CyQUANT™ LDH Cytotoxicity Assay (Thermo Fisher, C20300) and a FlexStation 3 plate reader using SoftMax Pro 7 (Molecular Devices).

### *In vitro* LPS-induced inflammation experiments

RAW 264.7 macrophages or BMDMs were seeded at a density of 250,000 cells per well into a 12-well tissue culture plate in 1 mL of media and left to incubate overnight. The indicated drug or vehicle control was added to each well, the plate was agitated and left to incubate for 10 minutes. The cells were then treated with LPS (1 µg/mL) or an equivalent volume of PBS, agitated, and left to incubate for 3 hours. Cells were washed with PBS, then RNA isolation was performed using RNeasy Plus Micro Kit (Qiagen, 74134). cDNA was produced using GoScript Reverse Transcriptase (Promega, A5004) and a quantitative PCR was performed using SensiFast SYBR (Bioline, 98020) and Bio-Rad CFX Connect thermocycler. Each datum represents an individual, biological replicate that is the mean of duplicate technical replicates. Fold-changes are all reported relative to untreated controls for each experiment and were calculated using the ΔΔCt method, with β_2_-microglobulin used as the housekeeping gene for each sample. The experiment was performed similarly with BV-2 microglia and alveolar macrophages, except that cells were plated at a density of 100,000 cells per well into a 24-well tissue culture plate with 1 mL of media per well.

### *In vivo* LPS-induced peritonitis mouse model

12-week-old, wild-type C57BL/6J mice were purchased from The Jackson Laboratory (Strain # 000664). An equal number of male and female mice were randomly assigned to each treatment group. All studies were conducted according to a protocol approved by the University of Virginia Animal Care and Use Committee (ACUC). Mice were socially housed with littermate controls and provided with free access to a standard chow diet and water. Baseline weights were taken <1 hour prior to the experiment for individual dose calculations. For the sub-lethal model, Mice were injected intraperitoneally with LPS (1 mg/kg), VPC01091.4 (30 mg/kg), both LPS and VPC01091.4, or vehicle alone (20% 2-Hydroxypropyl-β-cyclodextrin in PBS). The volume injected was 200 µL and solutions were sterile filtered (0.45 µm) prior to use. Mice were observed for 4 hours, with weight and symptoms assessed every 2 hours (t = 0, 2, and 4 hours). Mice were euthanized at 4 hours, then blood was collected via cardiac puncture and anticoagulated with K_2_EDTA. The inferior vena cava was severed immediately superior to the common iliac bifurcation, and the mice were perfused using 10 mL of ice-cold PBS into each cardiac ventricle. The lungs were dissected, coarsely homogenized, and partitioned into separate samples for RNA extraction and mass spectrometry analysis. Samples to be used for RNA were stored at −20°C in RNA*later*™ Stabilization Solution (Thermo Fisher AM7021) while samples for use in mass spectrometry were flash frozen in liquid nitrogen. Whole brains were physically dissected, coarsely homogenized, and stored for later processing as with the lung tissue. 10 µL of each whole blood sample was reserved for mass spectrometry analysis, 15 µL was used to obtain lymphocyte counts with a Hematology Analyzer (Heska Element HT5), and the remainder was used to produce plasma through centrifugation at 2,000 x *g* for 15 minutes at 4°C. Plasma cytokines were then measured using a multiplex mouse pro-inflammatory cytokine panel performed by the University of Virginia Flow Cytometry Core (RRID: SCR_017829). Cytokine values beyond the Limit of Detection (LOD) are reported as the LOD value. Tissue RNA was isolated from preserved samples using 30 minutes of automated homogenization at 30 Hz with 5 mm ball-bearings in RLT Plus Lysis Buffer (Qiagen TissueLyser II), RNA was then isolated using a RNeasy Plus Micro Kit (Qiagen, 74134). Whole-tissue cytokine expression changes were then measured using qRT-PCR as with *ex vivo* cell culture experiments. For the lethal dose model, mice were injected with LPS (25 mg/kg), both LPS and VPC01091.4, or vehicle alone (20% 2-Hydroxypropyl-β-cyclodextrin in PBS). Symptoms were scored according to a standardized clinical score sheet that assessed conjunctivitis, lethargy, degradation in hair grooming behavior, stool changes, and facial grimace (Table 1). Mice which lost more than 20% of their starting body weight or received a clinical score of 8 or greater were euthanized.

#### LC-MS/MS Sample Preparation

The sample preparation method was adapted from that published by Shaner *et* al (Shaner et al., 2009). Specifically, homogenized tissue (100 µL) was added to a mixture of methanol and chloroform (2 mL, 3:1, LCMS grade) and incubated at 48°C for 16 hours. The mixture was cooled to room temperature, an alcoholic KOH solution (200 µL, 1M in LCMS grade methanol) was added, and the mixture was further incubated at 37°C for 2 hours. The samples were neutralized with glacial acetic acid (20 µL) and centrifuged at 4°C for 10 min at 10,000 x *g*. The supernatant fluid was transferred to a glass vial and dried under a stream of nitrogen. The material was dissolved in LCMS grade methanol (500 µL), vortexed, and centrifuged at 4°C for 10 min at 10,000 x *g*. The supernatant fluid transferred to LC vials for analysis by LC-MS/MS.

#### LC-MS/MS Analysis

Analyses were performed using a tandem quadrupole mass spectrometer (Waters Xevo TQ-S micro) coupled to a UPLC (Waters Acquity h-class+) inlet equipped with a reverse phase C18 UPLC column (Waters BEH C-18 1.7 µm bead size, 2.1mm x 50mm). The method used is a modification of that published by Frej *et al (Frej et al., 2015)*. Specifically, the LC flow rate was set at 0.4 mL/min and the column temperature was 60^°C^. Mobile phase A consisted of water : methanol : formic acid (79:20:1) while mobile phase B was methanol : acetone : water : formic acid (68:29:2:1). The run began with 50:50 A:B for 0.5 min. Solvent B was then increased linearly to 100% B in 3.5 min and held at 100% B for 3 min. The column was re-equilibrated to 50:50 A:B for 1.5 min. A volume of 3 μL was injected on column. Both VPC01091.4 and FTY720 were analyzed in positive mode using MRM protocols as follows: VPC01091.4 (304.2 →269.2, voltages: cone 48, collision 12) and FTY720 (308.2→255.2, voltages: cone 4, collision 14). Quantification was accomplished using Waters TargetLynx ver. 1.4. The concentration of VPC01091.4 in tissues were calculated using the metric ratio to the internal standard (FTY720) and normalized to wet tissue weights.

#### Statistics and data processing

Data were analyzed using Microsoft Excel, Prism 9.5 (GraphPad Software) and Origin Pro 7.5 (OriginLab). Chemical structures were created using ChemDraw Professional 20.1.1 (PerkinElmer). Electrophysiology traces were plotted using Origin Pro 7.5, the remaining data were plotted and statistically tested using Prism. Outliers were determined using the Robust regression and Outlier removal (ROUT) function within Prism. IC_50_ values were determined by plotting the percentage of maximum current observed as a function of applied drug concentration. A non-linear regression was fit using a standard Hill slope of −1.0 and the model: *Y* = 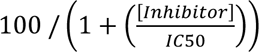. Where appropriate, data are presented as individual values, otherwise the sample sizes and precision measures are indicated in the figure legends. P values of less than 0.05 were considered statistically significant.

---

**Figure S1:**
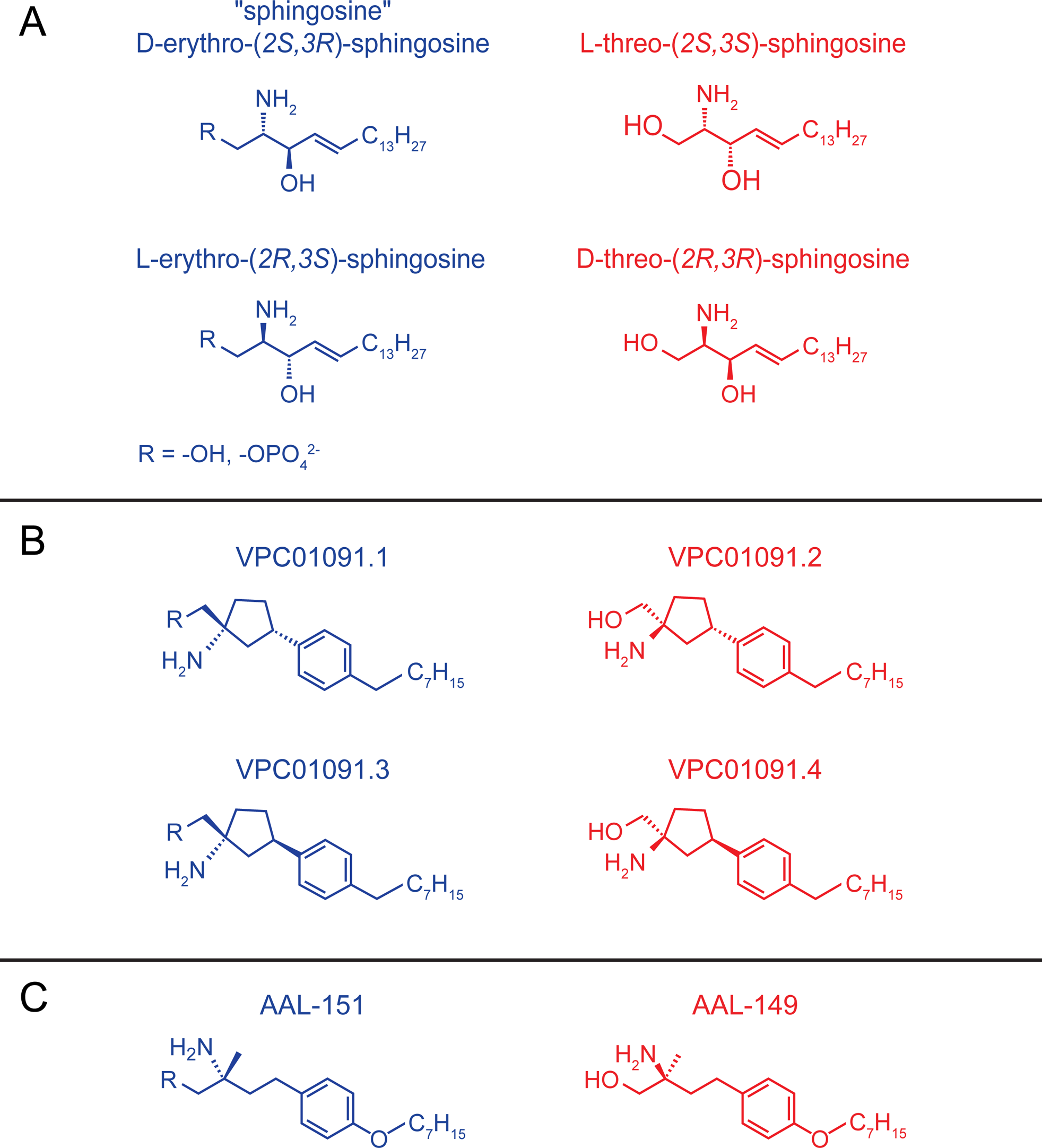
Stereoisomers of sphingosine, VPC01091, and AAL-149. (A) There are four stereoisomers of sphingosine, only D-erythro-sphingosine occurs naturally. The erythro-isomers are substrates for sphingosine kinases and are capable of being phosphorylated *in vivo* while the threo forms both act as competitive inhibitors of sphingosine kinases and are not appreciably phosphorylated *in vivo*. Phosphorylatable isomers are shown in blue. (B) The four diastereomers of VPC01091 are shown. VPC01091.1 and VPC01091.3 (blue) are substrates for sphingosine kinases, whereas VPC01091.2 and VPC01091.4 (red) are not. All four stereoisomers can be chemically phosphorylated, such as with VPC01091.2-phosphate used in the present study. (C) AAL-149 and its phosphorylatable enantiomer, AAL-151 are shown.

**Figure S2:**
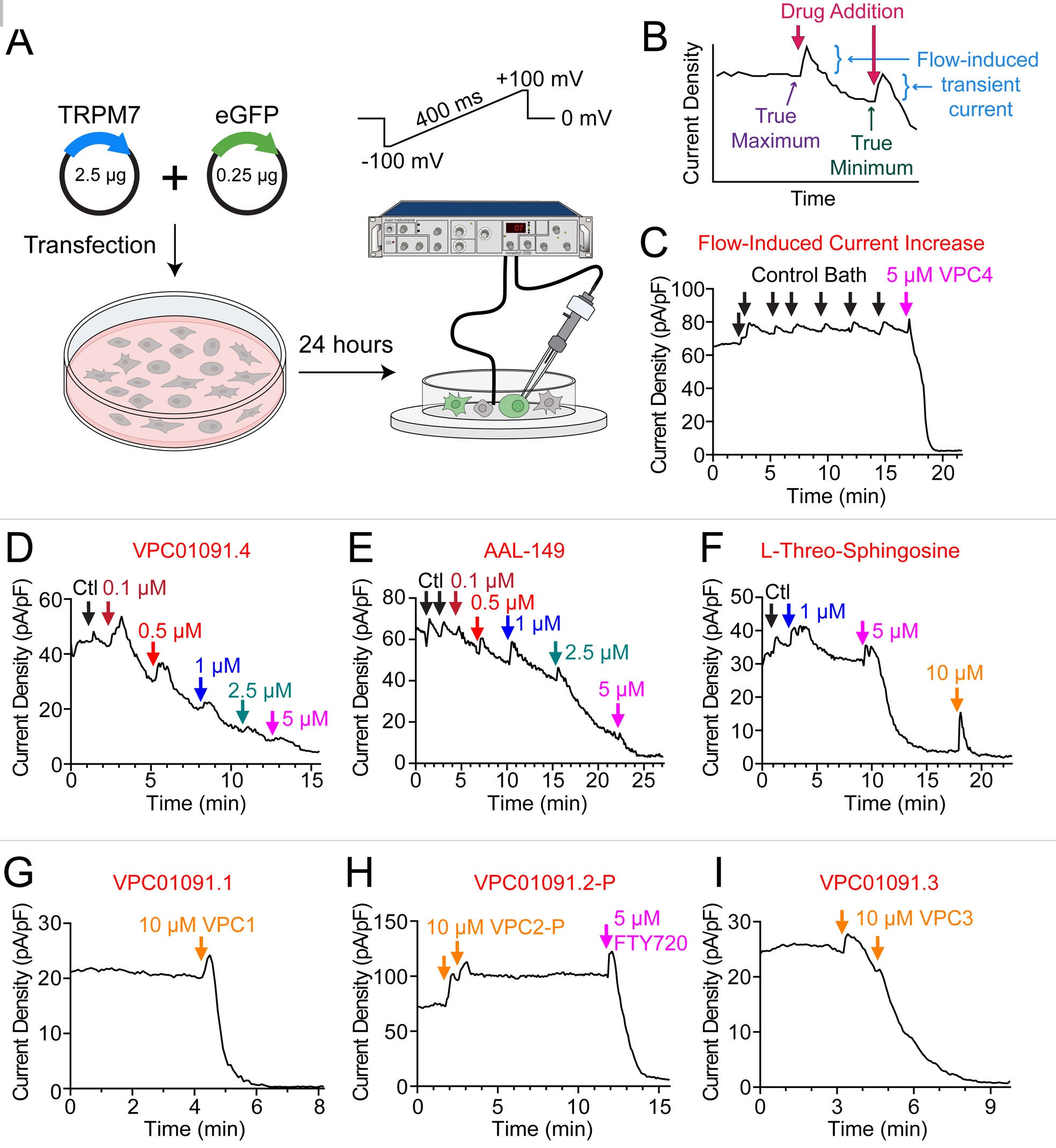
VPC01091.4, AAL-149, and L-threo-sphingosine inhibit TRPM7 current. (A) Schematic depicting the experimental setup for recording whole-cell currents of overexpressed mouse TRPM7 from HEK 293T cells. HEK 293T cells were transfected with a 10:1 ratio of WT mouse TRPM7 and eGFP using jetOptimus transfection reagent. After 24 hours, recordings were obtained from GFP-positive cells using an Axopatch 200b amplifier with a 400 ms ramp from −100 to +100 mV and a holding potential of 0 mV. (B) Schematic depicting the interpretation of peak current density at +100 mV vs time graphs. The perfusion of bath solution produces a flow-induced transient current increase (shown in blue). The current observed prior to the flow-induced transient is taken as the true maximum current. The trough observed after the decay of the flow-induced transient and prior to the next dose of inhibitory drug is taken as the true minimum current. The fraction of current remaining for each drug dose is taken as local minimum current over the true maximum current for the recording. (C) Example TRPM7 current density at +100 mV vs time graph for a recording in which control bath is repeatedly added to the recording chamber. Each addition of control bath produces a flow-induced transient current that decays back towards the pre-infusion maximum. Later addition of 5 µM VPC01091.4 leads to current inhibition. (D) Example TRPM7 current density at +100 mV vs time graph for increasing concentrations of VPC01091.4. Arrows depict the time at which the recording solution was replaced with control solution or VPC01091.4 at the indicated concentration. (E) Example TRPM7 current density at +100 mV vs time graph for increasing concentrations of AAL-149. (F) Example TRPM7 current density at +100 mV vs time graph for increasing concentrations of L-threo-sphingosine. (G – I) Example TRPM7 current density at +100 mV vs time graphs for VPC01091 stereoisomers. Application of 10 µM VPC01091.1 (G) or VPC01091.3 (I) leads to current inhibition. Application of 10 µM VPC01091.2-P (H) does not lead to current inhibition, but the current is later blocked through the addition of 5 µM FTY720.

